# Heritability jointly Explained by Host Genotype and Microbiome:Will Improve Traits Prediction?

**DOI:** 10.1101/2020.04.25.061226

**Authors:** Denis Awany, Emile R. Chimusa

## Abstract

As we observe the 70^*th*^ anniversary of the publication by Robertson that formalized the notion of ‘heritability’, geneticists remain puzzled by the problem of missing/hidden heritability, where heritability estimates from genome-wide association studies (GWAS) fall short of that from twin-based studies. Many possible explanations have been offered for this discrepancy, including existence of genetic variants poorly captured by existing arrays, dominance, epistasis, and unaccounted-for environmental factors; albeit these remain controversial. We believe a substantial part of this problem could be solved or better understood by incorporating the host’s microbiota information in the GWAS model for heritability estimation; ultimately also increasing human traits prediction for clinical utility. This is because, despite empirical observations such as (i) the intimate role of the microbiome in many complex human phenotypes, (ii) the overlap between genetic variants associated with both microbiome attributes and complex diseases, and (iii) the existence of heritable bacterial taxa, current GWAS models for heritability estimate do not take into account the contributory role of the microbiome. Furthermore, heritability estimate from twin-based studies does not discern microbiome component of the observed total phenotypic variance. Here, we summarize the concept of heritability in GWAS and microbiome-wide association studies (MWAS), focusing on its estimation, from a statistical genetics perspective. We then discuss a possible method to incorporate the microbiome in the estimation of heritability in host GWAS.

## 1 Introduction

Over a century ago, Weinberg [1], cognizant of the fact that phenotypic variation results from a combination of genetic and environmental factors, suggested methods of delineating genetic from environmental components of total phenotypic variability. This and subsequent works on statistical separation of the environmental and genetic variation in general populations, culminated to the specification of the fraction of the total phenotypic variance due to the genetic variance - a measure eventually termed ‘degree of heritability’ or simply ‘heritability’ in the genetic community [2]. A distinction is, however, necessary between *total* (or *broad sense*) and *additive* (or *narrow sense*) heritability. The former measures the full contribution of genes, which includes additive, dominance and epistasis components, while the latter captures only the additive contribution of genes to phenotypic variance.

It is now known that many common human diseases and traits are complex, resulting from the joint effect of host genetic and environmental factors. Indeed, genome-wide association studies (GWAS), which assays hundreds to millions of genetic markers - commonly single nucleotide polymorphisms (SNPs) - in thousands of individuals, have uncovered hundreds of genetic variants associated with many common polygenic inherited diseases and traits; revealing scores of previously unknown key biological pathways, and providing valuable insights into the complexities of their genetic architecture [3–5]. Despite this, however, GWAS has been puzzled by the apparent rather low proportion of the estimated heritability, which is far less than that obtained from familial studies - the difference being referred to as *missing/hidden* heritability. A classic, often cited, example is the human height where whereas the estimated heritability is 80%, the (narrow sense) heritability estimate with tens of thousands of people is only about 5% [6]. Many possible, and debated, explanations have been offered for this discrepancy, including sub-optimal sample size, poor detection of variants by genotyping arrays, dominance, epistasis, and shared environment [see references [4, 6] for excellent reviews]. While many investigators have argued that a considerable part of the missing/hidden heritability may be attributed to non-additive effects such as dominance and epistasis, several recent empirical studies have found no strong effects from them [7–9]. A similar observation has been made for epigenetic effects. While epigenetic variation, including methylation that has been suggested as another possible source of missing/hidden heritability, a recent study of body mass index found genetic predictors and methylation to be non-overlapping, suggesting the latter represented the environmental effects on this phenotype; for human height, methylation profiles did not explain any variation [9, 10]. Although the volume and scope of these studies are certainly not optimal, and hence the conclusions may not be entirely generalizable, they do corroborate the significant contribution of non-genetic factors to inflating heritability estimates from GWAS.

On the other hand, in parallel to GWAS, Microbiome-wide Association Studies (MWAS), have been successful in identifying bacterial taxa that are associated with a variety of conditions, such as obesity, major depression, colorectal cancer, and inflammatory bowel disease [11]. Interesting, however, recent twin-based studies have reported the heritability of the human microbiome. For example, in the largest twin cohort to date, Goodrich et. al. (2014) [12], using the gut microbiome samples, found a number of bacterial families to be heritable, with *Christensenellaceae* and having the highest heritability (*h*^2^ = 0.39). These interesting findings have raised enthusiasm, in as much as questions, among researchers on the implication of the microbiome on human health and the degree to which the human genotype versus the microbiome and the environment determines phenotypic variability.

Although GWAS and MWAS have been viewed as parallel fields, it has become increasingly apparent that time is ripe to shift away from the unidirectional host-centric and microbiome-centric interpretation to a more comprehensive view in which both host genetic and microbiome are considered as integral unit in analysis of phenotypic variability. The main reason for this is that if everything external to the human host is defined as the “environment”, then the environment in this case is another living organism. From ecological viewpoint, fluxes between biotic and abiotic components in an environment relies almost entirely on the abilities of the biotic components to extract and use the abiotic [13]. This is, however, not the case for the host’s microbiota, where this exchange is highly regulated by the host, for example through immune system [13, 14] and metabolic pathways [15, 16]. This associative trajectory, involving the host and microbes together with their collective genomes, greatly influences host biochemistry [16]; the ultimate result of which is the modulation of the host’s phenotypic expression. Thus, whether or not the host’s genome and microbial components both explain the same phenotypes, it is clear, from a statistical genetics perspective, that inclusion of both would improve statistical power to detect truly associated causative variants.

Apart from enabling the detection of causal variants, this comprehensive view, as also pointed out by other authors [13, 17], has the potential to narrow or provide insights into the missing/hidden heritability gap in GWAS for two reasons. First, while, by definition, heritability measures phenotypic variance attributable to genetic variance, GWAS only take into account the genetic variance in human cells and does not consider all the contributory role of the microbiome on the phenotype [13, 17]. Second, as a benchmark, the heritability estimates from familial studies, in which identity is inferred by kinship, are inflated because the observed phenotype is the resultant effect of the host’s genotype and microbiota (and of course, in addition to other external factors) [17]. Therefore, incorporating the host’s microbiota information with the genotypes will likely improve the estimates of contributors to heritability, and eventually facilitate the determination of either additional genetic variants to explain further proportions of heritability or the proportion of genetic variance that is already explainable by the already known variants. This add will additional anable the improvement of polygenic risk score for potential clinical utility.

In this review, we discuss the prospects for unifying the estimate of heritability expalned from GWAS and MWAS in uncovering the host’s genetic basis of human phenotypes. We focus on heritability estimation, from a statistical genetics perspective, and summarize the methodological approach to estimate (narrow sense) heritability. Finally, we suggest a method to incorporate the microbiome in the classical estimation of the heritability in human genome-wide association studies and conclude with a discussion of research areas where further work on both heritabilty and human traits prediction are needed.

## 2 Heritability in GWAS

Thousands of reports of GWAS mostly of European-ancestry encompassing larger samples, with some studies reaching up to million subjects have enabled the development of various heritability models to predict the genetic liability of human traits. Therefore, the clinical utility of the heritability has largely been explored in populations of European-ancestry and enabled applications in polygenic risk score and both genetics testing and counselling.

### 2.1 Broad-sense and narrow-sense heritability

Heritability is a measure of the relative contribution of genetics to a phenotypic expression. Its estimation centers around the measure of variability, which makes sense only if the phenotype is quantitative. For categorical phenotypes, therefore, one typically postulates it to be resulting from some underlying quantitative (continuous) variable, often called *liability*, which has a *threshold* that defines the intervals corresponding to the different states of the categorical variable [18]. The basic idea of heritability estimation is simple, at least in theory: partition the variance of a phenotype into components attributable to the different factors that are known to affect the phenotype, and determine the ratio of the genetic variance component (assuming genetics modulates the phenotype) to the phenotypic variance.

Suppose a quantitative trait is modulated by its overall genotype *G* and the exposure environment *E*, where *G* can be partitioned into additive 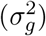, dominance 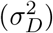, and interaction 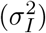 components.

That is,

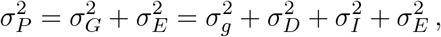

where 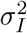 may be refer to additive-by-additive, dominance-by-dominance, additive-by-dominance as well as many higher-order interaction terms [19]. Broad sense heritability *H*^2^ is the ratio of the total 2 genetic variance to the phenotypic variance: 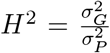, and expresses the degree to which genotype determine phenotype of individuals. Narrow sense heritability *h*^2^, on the other hand, is the ratio of the total genetic variance to the phenotypic variance: 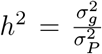, and expresses the extent to which individual’s phenotypes are determined by genes transmitted from parents. It therefore determines the degree of phenotypic resemblance between relatives, the observable genetic properties of a population and of the response of a population to external forces such as selection [20]. Broad sense heritability is of more theoretical interest than practical importance as it neither provide an understanding of the genetic properties of a population nor reveal the cause of phenotypic resemblance between relatives. Henceforth, as with all GWAS studies, we refer to narrow sense heritability (or simply heritability) in all subsequent discussions, unless stated otherwise.

### 2.2 Estimation of heritability in GWAS

We are typically interested in the *explained* heritability - defined as the ratio of heritability explained by a set of variants known definitely to be associated with the trait to the heritability explained by all genetic variants [those known (discovered) plus those not yet discovered] that are associated with the trait: 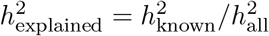; missing heritability being defined as 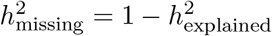 [4].

From a statistical genetics perspective, the additive variance 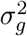 at a single locus is the genetic variance explained by the regression of the expected value of the phenotypic mean in each genotypic class on the genotype [7, 21], or put differently, heritability is the coefficient of the regression obtained from the regression of additive genetic effect on the phenotype. Therefore, as we detail below, 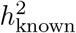 can be readily estimated from observed genotype-phenotype data using regression in a ‘bottom-up’ approach. The estimation of 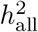 is, however, not straight forward because we do not know the complete repertoire of genetic variants associated with the trait; all we can do therefore is to infer it from phenotypic correlations obtained from population data in a ‘top-down’ approach.

Given a GWAS, let *y_i_, i* ∈ {1, ⋯, *n*} be the quantitative phenotypes measured on *n* individuals; *g_i_* = {*g*_*i*1_, ⋯, *g_im_*} the genotype of the *i^th^* individual for the *m* typed SNPs, with minor allele frequencies *P_j_, j* ∈ {1, ⋯, *m*}.

Employing the additive model, we have

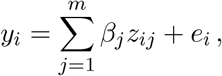

where 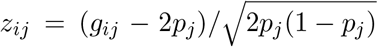 is the normalized genotype and *y* is normalized phenotype, having mean 0 and variance 1.

If *S* is the subset of statistically-associated (assumed here to be causal) variants obtained from the GWAS, then, as the phenotype is normalized (i.e. 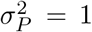 = 1), the additive variance is 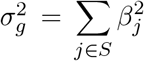 and the heritability is 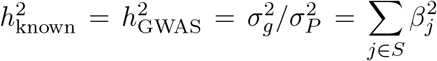, the sum of squared effect sizes for the normalized genotypes over the causal variants [4].

It is important to note, however, that the full set of causal variants are unknown; that is, the causal variants identified by the GWAS here is only a subset of causal variants. Consequently, 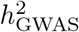 represents only the lower bound of the true heritability 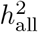. The difference between 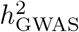 and 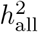 is termed the *missing* heritability of the phenotype. This difference can be attributed to several factors. First, the non-additive genetic variance such as epistasis that is not included in estimation of heritability; the presence of such non-additive variations have been shown to inflate heritability estimates [4, 22]. Second, the exclusion of causal variants due to, say stringent GWAS significance threshold or low effect sizes, for example, can lead to underestimation of the heritability. Likewise, false positive results would inflate observed estimates.

In practice, having estimated 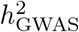, it is often of interest to know the proportion of the explained heritability. In other words, we would like to answer the following question: *what proportion of the heritability do all SNPs that contribute to the trait explain?* This question requires us to estimate 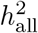.

The methodological estimation of 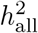 is a ‘top-down’ approach that hinges on recognizing the equivalence between then classical definition of heritability 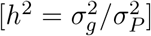 and the intuitive interpretation of the proportion of phenotypic variance explained by all causal variants 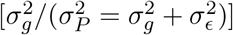.

If

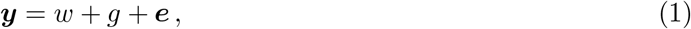

where *w* denote the fixed effects (including candidate SNP and optional covariates), *g* denote genetic effects assumed to subsume any genetic effects on the trait other than at the candidate SNP, 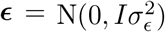; *I* being the identity matrix, then by treating *g* = ***Xβ*** as a random effect with 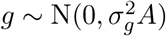, the variance-covariance matrix of ***y*** can be expressed as

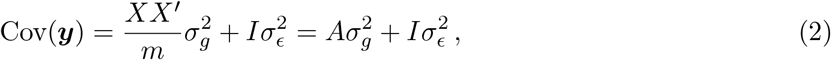

where *A* is the kinship (genetic relationship) matrix between pairs of individuals, defined over all causal loci [23], *m* is the number of causal variants, and 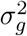 is the variance explained by the SNPs. Since *A* is defined over *all causal loci*, 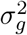 denotes the *variance explained by all causal SNPs*.

The parameters to be estimated are 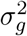 and 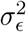, which can be done by using (2) in (1) and obtaining parameters optimization using the restricted maximum likelihood (REML) method. With these obtained, 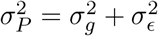 can subsequently be derived. In theory, this appears a trivial task. In practice, however, this is not the case because the estimation requires that we first obtain the matrix *A*, defined at the causal SNPs, but we do not know the causal SNPs. The traditional, and still used, approach involves using genetically related individuals from known pedigrees (family/twins) to estimate a kinship coefficient Φ, where, for example, Φ*_i_j__* is taken as 0.25 for siblings and 0.5 for twins; *A* is then taken to be equal to 2Φ [17, 24]. Clearly, this assumption does not necessarily hold since phenotypic resemblance may be influenced by other heritable factors, other than genotype; for example epigenetic modifications and the host’s microbiome. Indeed, the true covariance has been observed to vary around this assumed value [25]. Consequently, this can lead to inflation of the corresponding estimated heritability.

With the unveiling of genotype data, prodded by the advent of next generation sequencing, methods have been devised to estimate *A* from genotype data of unrelated individuals. This is done [26, 27] by postulating that the ungenotyped causal SNPs are tagged by the genotyped ones, and therefore although the set of causal variants is unknown, one can use all the SNPs genotyped in GWAS to estimate *A*, and use it as proxy for *A*_causal_. The key point to note here is that all genome-wide SNPs are used, not only the genome-wide significant SNPs. Indeed, this methodology proceeds without conducting any test of association between individual SNPs and the phenotype. The genetic relationship between individuals *i* and *j* is estimated using standardized genotype by

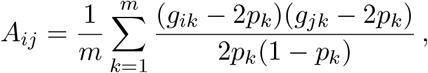

where *p_k_* is the allele frequency of the *k^th^* SNP. With *A*, defined at all causal SNPs, now obtained, (1) can be solved to estimate 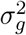 and 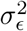, from where the variance heritability explained by all causal SNPs 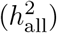 can be determined from 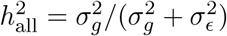.

In the estimation of both 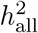 and 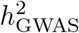 above, it is assumed that all causal SNPs have been genotyped or at least, are tagged by the genotyped SNPs. Accordingly, if the SNP array used does not fully cover the set of common genetic variants in the GWA study population, the resulting heritability estimates are likely to be smaller than the actual value, however large the sample size maybe.

Finally, it must be noted because the set of causal variants are unknown and we have to rely on SNPs being tagged by Linkage Disequilibrium (LD), and yet LD strength declines with increasing difference in minor allele frequency (MAF) between SNPs [28, 29], some causal variants in the low frequency spectrum may not be tagged; and, as a result will not play part in heritability estimates. Moreover, MAF of a disease allele can be population-specific [30, 31]. Because of this, the heritability estimate can be population specific. Indeed, it has recently been shown, for admixed population, that the narrow-sense heritability vary according to the local ancestry of the study population [32].

We point out that the general Linear Mixed effect Model (LMM) in Eq.(1) remains the fundamental method for heritability estimation; albeit, several different variants of it have been proposed in a bid to improve performance or allow estimation in different contexts; for example, to address environmental variations across samples [33], to refine the model for categorical traits [34], or to perform estimation in context-specific scenarios (such as genotype-environment, and genotype-sex contexts) [35]. Besides the REML-based methods implemented in LMMs, regression of phenotype correlations on genotype correlations (LDSC) and regression of phenotype correlations on genotype correlations (PCGC) are the other broad categories of statistical frameworks for heritability estimation (see **Box 2**). While each general method has its inherent strength and limitation, it is important to highlight the limitations to avoid potential pitfalls when applying a particular method. Currently implemented REML-based methods underestimates heritability in case-control studies [36–38], possibly due to due to case-control ascertainment biases [38]. LDSC methods can produce biased estimates in the presence of binary covariates with strong effects [36]. Meanwhile, the PCGC methods, although effective for case-control studies, can suffer from loss when there is ascertainment bias or when the genetic correlation is not constant (that is, inhomogeneous) across the allelic frequency spectrum at the risk loci. These being said, all methods suffer power loss when applied to cohorts from ancestrally divergent populations [36].

#### Box 2: Common Tools to Estimate Heritabilty

| Software Tool | Statistical Model | Trait | Note | Link |
| --- | --- | --- | --- | --- |
| GCTA | Linear Mixed Model (LMM) | Quantitative | Most routinely used tool for complex traits. Has been applied to a diverse array of traits, including human complex traits, transcriptional expression traits, and microbiome data. Generally effective for large sample size. | [27] |
| GxEMM | LMM | Quantitative and binary | Specifically designed for estimation in context-specific scenarios, including gene-environment interaction. Can accommodate modest sample sizes. | [35] |
| OpenMx | Structural Equation Modeling (SEM) | Quantitative and binary | Specific to classic twin-based study. Current implementation more appropriate for quantitative traits only. | [75] |
| GEAR | Phenotype correlations on genotype correlations regression | Quantitative and binary | The method originally proposed for linkage studies, achieves good performance with small samples. | [76] |
| MetaSex | LMM | Quantitative | Specifically designed to account for potential sex difference in genetic architectures. | [77] |
| StructLMM | Structured LMM | Quantitative | Purposely designed to model gene-environment interaction. | [78] |
| LDSC | LD score regression | Quantitative and binary | Estimating and partitions SNP heritability by functional annotations. Uses summary statistics as input data | [79] |
| SumHer | LMM | Quantitative and binary | Estimating and partitions SNP heritability by functional annotations. Uses summary statistics as input data. Key difference from LDSC is that it allows the user to specify the heritability model. | [80] |
| PCGC | Phenotype correlations on genotype correlations regression | Quantitative and binary | Performs better than GEAR in case-control studies. | [36] |
| LAMatrix | LMM | Quantitative | Estimates heritability by local ancestry, global ancestry, and degree of population differentiation at causal regulatory variants for gene-expression traits in admixed populations. | [81] |

### 2.3 *cis* and *trans* heritability of transcriptional regulation

While the pursuit for genetic variants underlying complex human diseases continues, results from the decade-long GWAS showed that over 90% of disease-associated variants lie in non-protein coding regions of the genome, for example in promoter regions, enhancers, and structural elements [39–43]. Since these non-coding DNA elements does bind proteins and RNA molecules which cooperate to regulate the function and expression of protein-coding genes [44], a prevailing hypothesis that human genetic variants impact traits via regulation of gene expression levels [40, 45–48]. This has motivated expression quantitative trait loci (eQTL) studies using genome-wide gene expression and genotype data, to explore the genetic basis of variability in gene expression; serving to potentially illuminate the bridge between statistical association and biological mechanism of a genetic variant on a phenotype. To this end, quantifying the heritability of gene expression is central to understanding its genetic basis and, ultimately, its contribution to host phenotypic diversity. Gene expression is known to be controlled by both *cis* eQTLs (defined as eQTLs located close, say within 250 kb - 1 Mb, to the gene it regulates) and *trans* eQTLs (defined as eQTLs located far, outside the cut-off distance, from the gene it regulates) [49], and therefore, when studying the heritability of gene expression, it is of biological interest [50] to express heritability in terms of *cis* heritability, 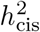, (heritability due to genetic component close to the regulated gene) and *trans* heritability, 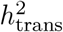, (heritability due to genetic component far from the regulated gene); so that the total heritability of a gene expression, 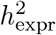, is given by 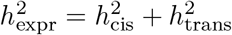. That said, *cis*-variation is often considered the primary driver of phenotypic variation; albeit, it is also more difficult to detect *trans*-acting eQTLs due to limitations in statistical power as their effect sizes are small [51, 52].

If we take the 250 kb ‘cis window’, for example, then the 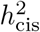 (narrow-sense) would be formally defined as the proportion of ‘gene expression phenotype’ explained by the additive effect of SNPs in a 250-kb window of the gene, whereas 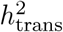 (narrow-sense) would be the proportion of ‘gene expression phenotype’ explained by the additive effects of SNPs outside a 250-kb window of the gene [53]. Methodologically, similar to the heritability estimation for complex traits, the heritability of gene expression can be estimated by two general approaches. The first involves using genetically-related individuals in the classical twin-based study design [54], where identity-by-descent (IBD) sharing across the genome is assumed to be 0.5 and 1 for dizygotic and monozygotic twins, respectively. The second alternative uses unrelated individuals where SNPs (measured plus tagged) within the *cis* window are used to define genetic-relatedness among individuals, and the estimation proceeds using the linear random effects model [27]; similar to classic heritability estimation of human complex traits. As the number of SNPs considered within *cis* window is small, 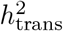 can be estimated with high precision [49, 55].

A number of recent studies have estimated heritability of gene expression across different human tissues. In a total of 856 female twins recruited from the TwinsUK resource, Grundberg and others [51] estimated 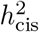 of gene expression for adipose, lymphoblastoid cell lines (LCLs) and skin tissues; obtaining, respectively, 26%, 21% and 16%. Importantly, having also estimated 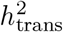, they reported that 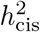 constituted between 30-36% of the total heritability, but up to 40% of 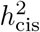 is missed when only common SNPs (MAF > 5%) are used in *cis* eQTL mapping. The important implication of this finding for host GWAS are that low frequency and rare variants may account for a substantial proportion of the unexplained *cis* heritability (for transcriptional regulation) and 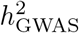 (for complex human traits), and that the action of host genetic polymorphisms on human diseases may be mediated by gene regulation, as the estimated 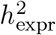 is enriched in genes previously identified via GWAS in a broad range of diseases. In another recent study to characterize the genetic basis of human gene expression [53], narrow-sense *cis* heritability of *LCL* gene expression was estimated to be approximately 8.2%. The authors found that singletons accounted for the vast majority (25% compared with al other MAF bins) of this heritability, and over 90% of this was due to alleles of ultra-low frequencies (MAF< 0.01%). Taken together, these findings suggest much of the missing (or unexplained) heritability of complex traits may be due to variants in the low-frequency spectrum, and transcriptional regulation represent at least one intermediary bridge between host genotype and phenotype.

## 3 Heritability in MWAS

Although the role of microbes in health and physiology has been known for over a century, only recently have the roles of these microbes together with their collective genome - the microbiome - in the pathogenesis of many common human diseases and traits become apparent, through microbiome-wide association studies (MWAS). MWAS in which the compositional and functional diversity of the microbiome is assessed at various taxonomic ranks (e.g species or genus level) in tens or hundreds individuals, represent a powerful new tool for investigating the microbiome basis of complex traits and diseases. To date, these studies have identified several microbiome-disease/trait associations [56]. The success of GWAS provided an optimistic outlook for MWAS and the observation of host genotype-microbiome interaction led to works on the heritability of the human microbiome.

Recent studies have shown that the microbiota is vertically transmitted from mother to offspring; albeit, the role, importance, and transmission mode of prenatal microbial colonization are still unclear [57, 57–59]. However, extensive colonization begins postpartum [57, 60]. Vertical transmission via breast milk, and horizontal transmission through factors such as mode of delivery (vaginal or caesarean section), feeding method (formula or breastfeeding), and social interactions are among the crucial factors in the development of the infant microbiome [57, 60, 61]. The transmission of the microbiota across humans is corroborated by the congruence of phylogenetic tree of intestinal bacterial microbiota and humans [62, 63]. Since microbial information can be transferred to offsprings and microbes have co-evolved with their human host for millions of years, it is reasonable to expect the former to hold information on latter’s phenotypic plasticity [62].

To estimate heritability of the microbiome, one can apply the standard Additive Genetics, Common Environment, Unique Environment (ACE) model, treating the abundance of each human-associated microbe as a quantitative trait. Heritability in then is estimated by determining variation in microbial taxon abundances (as measured by within-community alpha diversity measures such as observed species and Shannon diversity or by between-communities beta diversity measures such as UniFrac and Bray-Curtis metrics) that is attributable to human genetics. To date, twin studies invoking Falconer’s formula

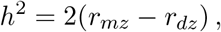

where *r_m_* and *r_d_* are the correlation between pairs of mono-zygotic and di-zygotic twins respectively, has been the basis of heritability calculation [12, 64]. These studies have provided clues into the nature and extent of host-microbiome association: bacterial taxa observed to be consistently heritable include *Christensenellaceae, Actinobacteria, Firmicutes*, and *Tenericutes*, while *Bacteroidetes* phylum were generally not heritable [12, 64]. It is important to note, however, that the volume of such research is still small and further data will lend insight into this link.

## 4 Incorporating the microbiome in heritability estimation

It is now known that the majority of the common complex phenotypes are the result of the contributory role of host genetics, the microbiome, and other environmental factors. How these components do combine to determine phenotypic expression is certainly unknown. The simplest model is to assume either the contribution of the environment and the genetic variants that act additively or the environment and the additive effect of the microbiome (see *Box 1*). However, multiple lines of evidence suggest that host genetics and the microbiome do not act independently to shape observed phenotype [56, 65]. Indeed, several recent studies have reported weak effect of host genetics on the microbiome, both host genetics and microbiome have been independently implicated in the etiology of the same diseases/traits. Therefore, as also noted by [17], it would be useful to integrate host genetics and the microbiome in the same analytical model. The classical definition of heritability is limited. Although host genetics do contribute to phenotypic expression, the definition is based on the premise that host genetics and the environment as an integral component do have a contributory role on the phenotype. In light of host-microbiome symbiosis, it is pertinent that the host genetics and microbiome be viewed as a single unit representing ‘host community genotype’. Indeed, the shift towards the view of organisms as an ecosystems has been advocated [17] (see Figure 1).

### Box 1: Statistical Models for Heritability of Quantitative Human Traits

Quantitative genetics theory is traditionally developed for quantitative traits. Nevertheless, the theory can still be applied to a categorical trait by assuming it to be governed by some underlying quantitative latent variable, often called a liability, whose thresholds delimit the categories [18]. Let *Y* denote the random variable for the quantitative trait, and suppose *G, M* and *E* are the random variables for genotype, microbiome and environment respectively.

#### Model 1: Host’s Phenotype is Influenced by the Host’s Genotype and Environment

If we consider Model 1 and assume additive genetic effects and ignore dominance and epistasis effects, then the overall phenotypic variance can be decomposed as

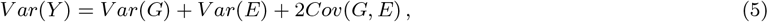

where *Var*(·) is the variance of (·), and *Cov*(·, ·) is the covariance of (·, ·).

In practice in heritability estimation, it is implicitly assumed that *G* and *E* are independent, that is, *Cov*(·, ·) = 0. The narrow-sense heritability of the host’s phenotype, 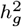, is then defined by

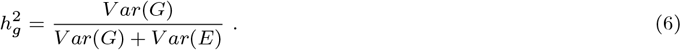

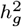 is the proportion of phenotypic variance attributable to genetic variance in the host.

#### Model 2: Host’s Phenotype is Influenced by the Host’s Microbiome and Environment

For this model, if we similarly assume additive effects of bacterial taxa on phenotype, and ignore between-taxa, and taxa-environment interactions, then, as above, the narrow-sense heritability of the host’s phenotype, 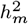, is then defined by

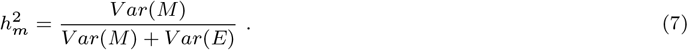

Analogous to 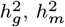 is the proportion of phenotypic variance attributable to variability in the host’s microbiome.

#### Model 3: Host’s Phenotype is Influenced by the Host’s Genotype, Microbiome and Environment

A perhaps more realistic model would be as in Model 3. In this case, assuming additive effects of SNPs and microbiome, we have

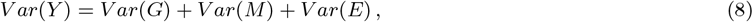

where we have imposed the assumption that *Cov*(*G, E*) = *Cov*(*M, E*) = *Cov*(*G, M*) = 0.

There are two possible heritability measures of interest that can be defined from this model, each with a different interpretation. First, is the ‘geno’ heritability which is the proportion of phenotypic variance explained by host genetic variance. That is,

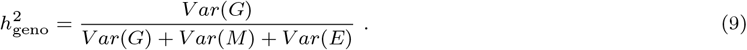

In other words, geno-heritability is the narrow-sense host’s genetic heritability of the trait. As with the classical interpretation of heritability, 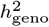 is taken to be obtained after accounting for the factors known to modulate the phenotype; the factors, in this case, being host’s genotype, microbiome and environment.

Second, in light of the holobiont theory of humans host and its microbiome [13], one can define ‘genobiome’ heritability as the proportion of phenotypic variance due to both host genetics and microbiome variances. That is,

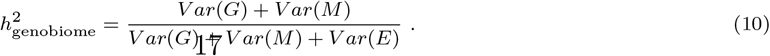

#### Methodological Implementation

Estimation of heritability can be performed by fitting a linear mixed model (LMM) or Haseman-Elston (H-E) regression. The LMM has been the standard tool for heritability estimation for various host quantitative traits. It has also recently been applied to heritability estimation for transcriptional expression traits, including gene expression, methylation level, and other molecular traits [71, 72]. In the standard LMM implementation, the phenotype of each individual is modeled as the sum of two sets of random effects; one based on the covariate(s) of interest (e.g individual’s genotype, or microbiome) and one based on environmental factors (see Eq.(1) in Main text for more detail). The parameters of the model are typically then fitted by maximizing the restricted maximum likelihood (REML) of the data [27], from where the desired heritability can be calculated from the estimated variance parameters.

Alternatively, especially when the sample size is small, as often seen with transcriptional expression traits, the H-E regression [36, 73] may be opted for, given its robustness for small sample sizes [53]. The idea here is to regress the phenotypic covariance on the genotypic covariance so that the resulting effect sizes, which will actually be the variance components, can be used to obtain the heritability of the phenotype under consideration. Again, in the practical implementation of H-E regression with transcriptional data, the SNPs are usually partitioned into *K* disjoint subsets based on minor allele frequency (MAF). The overall heritability of the trait is then the sum of heritability due to SNPs from each partition; this estimation, after partitioning SNPs, has been shown to correct for over- or under-estimation of heritability [53, 74].

Typically, in the transcriptional expression trait, the phenotypic covariance (denote it by, say, *Y*) is taken to be the upper triangle of the outer product of quantile-normalized log2(FPKM) [FPKM; Fragments Per Kilobase of transcript per Million mapped reads], and genotypic covariance (denote it by, say, *X*) defined as the upper triangle of a genomic-relationship matrix generated from all SNPs in the partition. For the *k*^th^ partition, 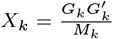, where *G_k_* and *M_k_* are, respectively, the standardized genotype and number of SNPs in the *k*^th^ partition. The mapping is then carried out with the usual linear regression; viz

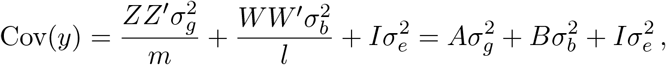

In this regression, the effect size for the *k*^th^ SNP partition represents the genetic variance of that partition; that is, 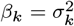. Thus, the total genetic variance due to all SNPs is 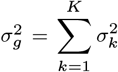. As the phenotype is normalized to unit variance, the (narrow-sense) heritability of the transcriptional expression trait, *h*^2^, is then equal to 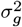.

**Figure 1:**
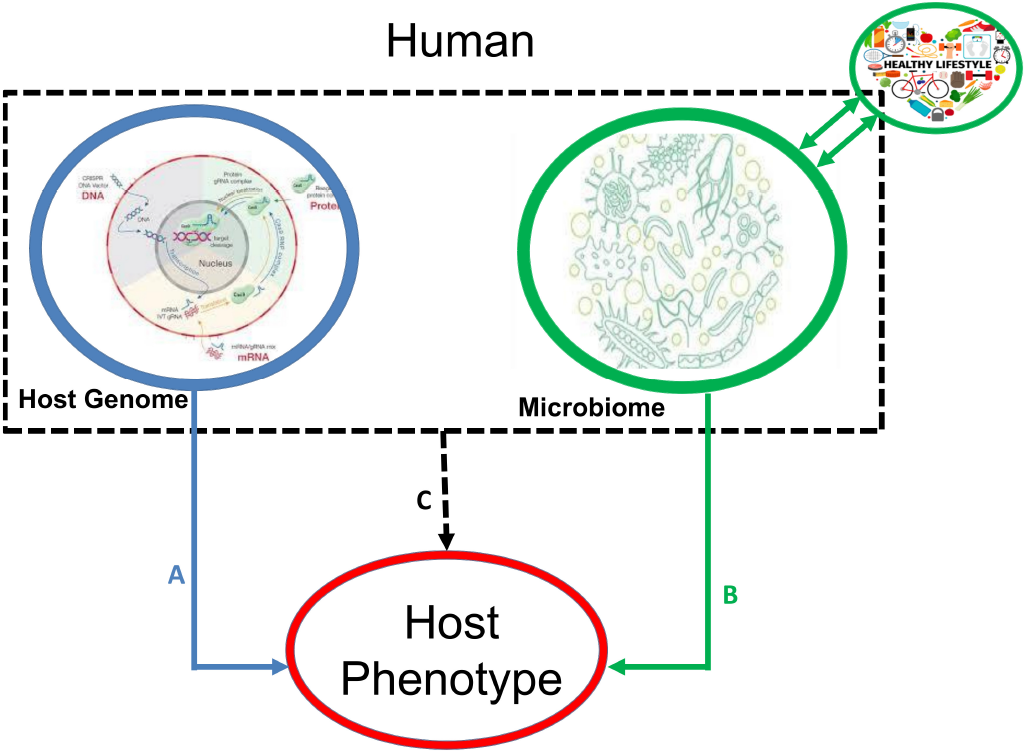
Conceptualization of host genetics and microbiome in heritability estimation for host phenotype. A: genetic heritability from host GWAS; B: microbiome heritability from MWAS; and C: heritability jointly explained by host genetics and microbiome.

In this community view, the microbiome can be integrated at various levels; for example, species, transcripts, metabolites, proteins, genes or their functional diversity. More generally, assume the phenotype (*y*) is contributory role of host genetics (*g*), the microbiome (*b*), and the environment (*e*). The model becomes

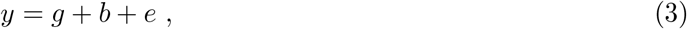

where 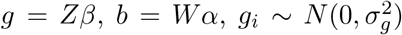, and 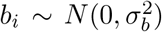. Whilst it would be possible to model interactions effects, we concentrate on the main effects part of the model, partly for simplicity of exposition as it is very difficult to identify interactions terms with a reasonable accuracy in such high dimensional setting [66–68] and partly because the first order of Taylor expansion of the model function can accurately approximate interaction effects, which are essentially encoded in lower order terms [68]. The phenotypic variance-covariance matrix is expressed as

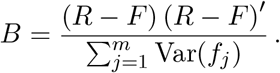

where *A* is a host genetic relationship matrix defined on the causal variants, *B* is the the microbial taxa similarity matrix defined over the associated taxa, 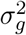 is the total total additive genetic effect and 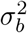 is the total total additive microbial effect, *m* and *l* is the total number of causal host genetic variants and microbial taxa, respectively.

Adapting the classical GWAS methodology, *A* can be estimated using the all SNPs genotyped in the study, as described above. The microbiome similarity matrix can, however, be defined in two ways. First, phylogenetic distance measures, which accounts for the phylogenetic relationship among microbial taxa, could be used to define sample similarity matrix *B.* This approach, that has a solid foundation in the field of microbial ecology, would allow *B* to incorporate the degree of divergence between sequences, thereby estimating similarity among individuals based on phylogenetic relatedness of microbial communities in their bodies. This idea is further supported by the observation that host humans have co-evolved with their microbiome [13, 63, 69] and microbiome-related phenotypes can be transmitted between phylogenetically close humans [13]. Second, for each microbial specie, the abundance data can be discretized into ‘categorie’ such as 0,1,2 corresponding to low, medium, high abundance respectively, based on some biologically-plausible scale. The frequency of each category can be calculated based on population values. This is, however, not straightforward as it involve knowledge of community-composition of each microbial specie. In the absence of this information, the complete set of individuals in the study samples may be used as proxy for the population. Once this information has been obtained, the sample similarity between the individuals can be obtained as follows:

Consider model (3) and let *R* be an incidence matrix that maps different categories of microbial taxa to each subject. In the above case, the elements of *B* are 0, 1, or 2. Following the usual definition of Euclidean distance similarity, if we *K* = *RR*′ then the diagonals of *K* give the subject’s relationship to itself while the off-diagonal elements gives the number of elements shared by the subjects. We are interested in investigating over-representation of microbial taxa. Accordingly, as in GWAS, it is possible to define a ‘reference category’ and determine the corresponding frequency. For our example, if we suppose category “2” is the reference category, then define *F* to be a matrix containing the frequencies of each category. The *j^th^* column of *F* is 1*f_j_*, where *f_j_* is the expected value of frequency of the reference category in the *j^th^* taxon. With this, the matrix (*R* − *F*) becomes the mean-centered form of *R*, which can essentially be interpreted as setting the mean value of taxa effects to zero. The microbial relationship matrix *B* is then calculated using

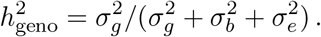

The normalization by 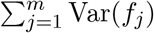 scales *B* in a way similar to the usual kinship matrix in GWAS. Finally, the phenotypic variance explained by additive variation at all common SNPs, which we denote by 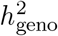, is calculated from

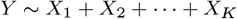

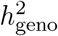 is the narrow-sense host’s genetic heritability of the trait, after accounting for the additive effects of host genetics and the microbiome. We call this ‘geno’ heritability.

Given that the microbiome co-evolved with its human host for millions of years, an emerging view is that of the ‘holobiont’ in which the human host and its microbiome is regarded as a single entity [13]. In light of this, one can define ‘genobiome’ heritability as the proportion of phenotypic variance due to both host genetics and microbiome variances. That is,

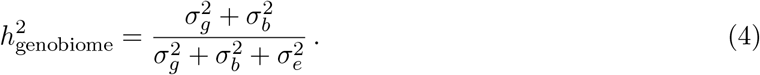

The genobiome-heritability, so defined, is the narrow-sense host’s genetic and microbiome heritability of the trait, and represents the heritability of the trait jointly explained by the host’s genetics and microbiome.

## 5 Concluding Remarks

Predicting the heritability of human traits is one of the critical goals in biomedical research and precision medicine. Today, thousands of reports of GWAS, encompassing larger samples mostly generated European ancestry. These efforts enabled the development of various models of heritability to predict the genetic liability of human traits. However, current heritability models developed using large-scale European-ancestry genomic data still misestimate the predictive power of heritability of most human traits and, they additionally suffer power loss when applied to cohorts from ancestrally divergent populations [36]. The clinical utility of predictive heritability of traits is still in its infancy stage and its application in genetics testing and counselling in real-world clinical populations is limited. Due to the differences in disease/traits prevalence, linkage disequilibrium (LD), genetics ancestry, environmental factors, microbiome profiles, causal or marginal effect sizes and, epistatic or gene-environment interactions between populations, heritability of trait derived from GWAS of European-ancestry samples can potentially misestimate the predictive risk power when applied to non-European populations [10]. In addition, most of non-European populations such as Africans exhibit significantly higher risk allele frequencies, of which ancestral risk alleles is higher than derived risk alleles commonly observed to populations of European-ancestry [5, 9], therefore new heritability approaches that leverage population-specific characteristic including epigenetics, genetics ancestry, host-genetics interaction with microbiomes are needed to improve the predictive power of the heritability of human traits.

The age of the microbiome is upon us, and the invaluable potential of the microbiome for host GWAS cannot not be overstated. Classical GWAS is based on the premise that the environment and disease are homogeneous among the study subjects. Insights gained from genetic and microbial epidemiological studies make it clear that this assumption does not generally hold, and can consequently reduce the power to detect truly associated causative variants. To this end, it is crucial that GWAS leverages the deterministic and stochastic factors that have known contributory role to phenotypic variability. In particular, given the association of host genetics and microbiome with the same phenotypes, the overlap of host genetic variants associated with the same traits, and the fact that the microbiome, unlike other abiotic environmental factors, is heritable and its variability has a genetic basis [70], it is pertinent that the microbiome be viewed as an integral part of the host rather than an external environmental factor. In this framework, the association mapping is performed on the host community, comprised of host genotype and its microbial community.

Moving forward, considering the additive effects of the microbiome in heritability calculation will be worthwhile as we seek to explain the ‘dark matter’ of missing/hidden heritability. Narrowing the missing/hidden heritability gap is of more than just an academic interest: knowing the heritability of a phenotype provides geneticists with the upper limit of the degree with which a phenotype can be predicted by identified variants. This is inevitable if we are to illuminate the dark path from genome-wide significant association to biological and medical application.

Beyond the missing heritability esoteric, the delineation of heritability in association mapping will be key in bridging the gap between statistical association and clinical translation in two broad ways. First, by quantifying the variance attributable to host genetics and microbiome, it will expand our understanding of complex disease architecture, which, ultimately, would guide design of experiments to fully dissect genetic and/or microbiome basis of disease aetiology. Second, is disease population risk stratification. With knowledge of the upper limit of risk stratification, disease risk models can be used to predict population-level risk of disease. The immediate benefit of this would be improved diagnosis, risk stratification, and disease management.

### Key Points

1. Heritability estimates from human genome-wide association study (GWAS) and microbiome-wide association study (MWAS) provide, respectively, the extent of host genetics and microbiome contributions to host phenotype.
2. The involvement of the microbiome on host phenotypes makes it apparent that the microbiome be integrated with host genotype in host trait association mapping.
3. A substantial portion of the unexplained (aka missing) heritability in GWAS could be accounted for if the microbiome variation is taken into consideration.
4. In light of the holobiont theory, the narrow-sense heritability jointly explained by host genetics and microbiome can be determined.
5. The clinical utility of heritability estimates, which include traits prediction, disease risk stratification and characterization of disease architecture, necessitates its pursuit.

## Acknowledgments

The author thank Delesa Damena and Imane Allali for helpful comments in an early draft of the paper. The authors thank CHPC (https://www.chpc.ac.za/) for providing computing facility.

